# A molecular phenotypic map of Malignant Pleural Mesothelioma

**DOI:** 10.1101/2022.07.06.499003

**Authors:** Alex Di Genova, Lise Mangiante, Alexandra Sexton-Oates, Catherine Voegele, Lynnette Fernandez-Cuesta, Nicolas Alcala, Matthieu Foll

## Abstract

**Background:** Malignant Pleural Mesothelioma (MPM) is a rare understudied cancer associated with exposure to asbestos. So far, MPM patients have benefited marginally from the genomics medicine revolution due to the limited size or breadth of existing molecular studies. In the context of the MESOMICS project, we have performed the most comprehensive molecular characterization of MPM to date, with the underlying dataset made of the largest whole genome sequencing series yet reported, together with transcriptome sequencing and methylation arrays for 120 MPM patients.

**Results:** We first provide comprehensive quality controls for all samples, of both raw and processed data. Due to the difficulty in collecting specimens from such rare tumors, a part of the cohort does not include matched normal material. We provide a detailed analysis of data processing of these tumor-only samples, showing that all somatic alteration calls match very stringent criteria of precision and recall. Finally, integrating our data with previously published multi-omic MPM datasets (*n*=*374* in total), we provide an extensive molecular phenotype map of MPM based on the multi-task theory. The generated map can be interactively explored and interrogated on the UCSC TumorMap portal (https://tumormap.ucsc.edu/?bookmark=746c4bc0e8bc4eb5f280cdd8lc7dcc783955faf2e2b493d0d205b7dle92b98c4).

**Conclusions:** This new high quality MPM multi-omics dataset, together with the state-of-art bioinformatics and interactive visualization tools we provide, will support the development of precision medicine in MPM that is particularly challenging to implement in rare cancers due to limited molecular studies.

## Context

Malignant Pleural Mesothelioma (MPM) is a deadly pleural cancer with currently limited therapeutic opportunities that translate into poor outcomes for patients. The latest WHO classification [1] recognises three different histopathological types, namely epithelioid (MME, median overall survival of 14.4 months), biphasic (MMB, 9.5 months), and sarcomatoid (MMS, 5.3 months). Multi-omic sequencing data [2,3] have been key in the identification of driver genes, developing and refining the characterisation of molecular profiles from initial discrete clusters to a continuum [4–6], and uncovering rare genotypes such as near-haploid genomes. Such advances have revealed the rich molecular heterogeneity in MPM, and have fueled the implementation of drug trials for more tailored MPM treatments. Despite their important findings, these multi-omic studies have profiled only a reduced representation of the MPM genome (primarily exomes) and have mainly focused on describing simple mutational processes (i.e. copy number alterations and point mutations). Therefore, there is still a need for comprehensive multi-omic datasets including whole MPM genome sequences to allow the study of complex mutational processes–e.g., whole-genome doubling (WGD), chromothripsis, extrachromosomal DNA (ecDNA)–that have been described in other cancer types [7–9] but not in MPM. Furthermore, understanding how genomic events impact tumor phenotypes remains poorly studied in MPM. Finally, given that MPM is a rare disease, the integration of different multi-omic studies is essential for reaching the statistical power needed to derive insightful biological conclusions from complex multi-omic datasets.

## Data Description

Here we describe the dataset generated by the MESOMICS project that collected more than one hundred MPM tumors with extensive clinical, epidemiological, and morphological annotations, and profiled their genome, transcriptome, and epigenome. Notably, MESOMICS prioritized the sequencing of whole MPM genomes rather than exomes, resulting in the largest set of MPM genome sequences available to date. This dataset has been deposited at the EMBL-EBI European Genome-phenome Archive (EGA accession No. EGAS00001004812). Additionally, we provide a comprehensive description of data quality control, and links to all bioinformatic pipelines used in the project, including state-of-the-art methodology for mutational calling in tumor-only specimens. Finally, in order to maximize the reuse potential of our MESOMICS data, we integrate our cohort with the previous multi-omic studies from Bueno *et al*. [2] and Hmeljak *et al*. [3] to generate the first multi-cohort molecular phenotypic map for MPM based on the multi-task Pareto optimum theory [10]. This interactive map provides a user-friendly way to explore the molecular data and to generate new hypotheses through custom statistical tests, based on the UCSC TumorMap portal [11]. The integrated and harmonized dataset resulting from these studies is available on GitHub [12].

Primary tumor specimens were collected from surgically resected MPM. As described in Mangiante *et al*. 2021 [13], among them 13 had two tumor specimens collected to study intratumoral heterogeneity; we report quality controls for all samples including these 13 additional samples, but only the piece with the highest tumor content as estimated by pathological review was selected for subsequent analyses, except for analyses that specifically focused on intra-tumor heterogeneity. The samples used in this study belong to the French MESOBANK. Our pathologist (FGS) classified all tumors following the latest WHO guidance and DNA, and RNA extraction methods are described in the methods section of our recent study [13].

### Quality control of omic data

#### Whole-Genome sequencing (WGS)

Whole-genome sequencing was performed by the Centre National de Recherche en Génomique Humaine (CNRGH, Institut de Biologie François Jacob, CEA, Evry, France) on 130 fresh frozen MPMs, plus 54 matched-normal tissue or blood samples (matched non-neoplastic tissue was not available for the other specimens). The Illumina TruSeq DNA PCR-Free Library Preparation Kit was used for library preparation and the HiSeqX5 platform from Illumina for the sequencing as described in [13]. The raw WGS reads were scanned by the FastQC software (v.0.11.5; using our nextflow pipeline https://github.com/IARCbioinfo/fastqc-nf) to determine the reads base quality, adapter content and duplication levels. The software MultiQC (v0.9) was then used to aggregate all the FastQC reports across samples.

The target read output for matched-normal tissue or blood (hereinafter called “matched-normal”) and for tumor tissues without matched-normal sample (hereinafter called “tumor-only”) was 900M reads (~30X genome coverage, **Figure 1A**). 1800M (~60X genome coverage, **Figure 1A**) were expected for tumor tissues with matched-normal samples (two sequencing lanes, hereinafter called “tumor-matched”). Overall, the median number of reads obtained approached or exceeded the target read output, with median and standard deviation by sample type equal to: matched-normal 889±50, matched-tumor 1786±163, and tumor-only 853±51 million reads (**Figure 1A**).

**Figure 1.**
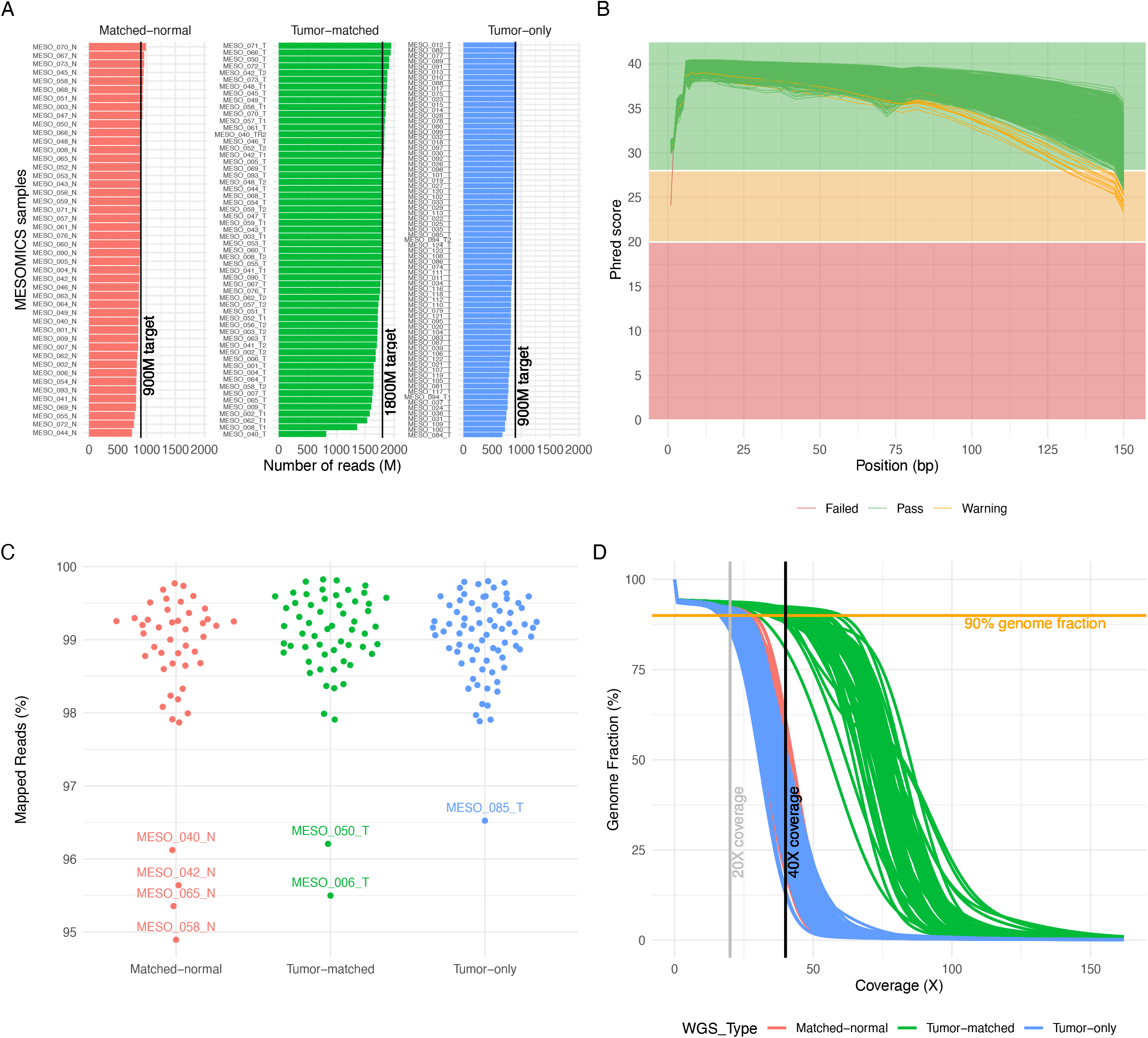
Quality control of Whole-Genome Sequencing (WGS) data. A) Number of reads per WGS type. B) Mean sequence quality score as a function of the position in the read in base pairs. Green lines correspond to files that passed the most stringent QC filters of software FastQC; orange lines correspond to files that passed a less stringent filter; and red to files that did not pass the filters. C) Percentage of aligned reads to the reference human genome. D) Cumulative genome fraction computed directly from the BAM files.

All samples displayed the expected mean quality score (30Q > 85% of bases) across all base positions of the read (**Figure 1B**). One exception is the MESO_050_N (a matched-normal sample) that on average had a good sequence quality score (**Figure 1B**) but displayed a low mean quality score for the first nucleotide of the read (24.08 Phred), which FastQC reported as a warning in the mean quality score module (**Figure 1B**). In fact, the FastQC report for this sample indicated that 25.17% of bases were not called at the first nucleotide of the read, suggesting that the base-calling process struggled in interpreting the DNA bases at this position and put an N instead. However, the reverse pair-end file of this sample had the expected sequence quality score over all the read positions (**Figure 1B**) and we decided then to include this sample in the subsequent analyses. The adapter content was lower than 1% for all sequenced samples (maximum 0.87% of total reads). The relative level of duplication found for every sequence per sample was on average 10.3% (min: 0% and max: 18.2%), this low level of duplication indicates that the prepared genomic libraries were diverse and likely covered a high proportion of the human genome.

Paired-end read mapping was performed with our nextflow [14] pipeline alignment-nf (v1.0, https://github.com/IARCbioinfo/alignment-nf). This pipeline includes the software qualimap (v2.2.2b) and MultiQC to generate comprehensive QC statistics reports from the WGS alignment files. The mean percentage of aligned reads was 98.93±0.81% (**Figure 1C**). The matched-normal and tumor-only samples displayed a mean genome coverage higher than 30X (**Figure 1D**). The matched-tumor displayed a mean genome coverage of 60X (**Figure 1D**). Finally, 90% of the reference genome was covered by at least 22, 20, and 43 reads for matched-normal, tumor-only, and matched-tumor samples, respectively (**Figure 1D**).

#### RNA Sequencing data

RNA sequencing was performed on 126 fresh frozen MPM in the Cologne Center for Genomics. Libraries were prepared using the Illumina^®^ TruSeq^®^ RNA sample preparation Kit, the Illumina TruSeq PE Cluster Kit v3, and an Illumina TruSeq SBS Kit v3-HS subsequent sequencing was carried out in an Illumina HiSeq 2000 sequencer, as described in [13].

The resulting raw reads files were processed using our nextflow RNA-seq processing pipeline (https://github.com/IARCbioinfo/RNAseq-nf v2.3), as described previously [13,15], that performs reads trimming (Trim Galore v0.6.5), and mapping to reference genome GRCh38 (gencode version 33) with STAR (v2.7.3a) [16]. We also improve the alignments as described previously by performing assembly based realignment (nextflow pipeline https://github.com/IARCbioinfo/abra-nf, v3.0), using software ABRA2 [17] and base quality score recalibration (nextflow pipeline https://github.com/IARCbioinfo/BQSR-nf, v1.1), using GATK v4.1.7.0 [18]. Gene-level quantification was performed using software StringTie (v2.1.2) (nextflow pipeline https://github.com/IARCbioinfo/RNAseq-transcript-nf, v2.2). Quality control of the samples was performed using FastQC (v0.11.9, https://www.bioinformatics.babraham.ac.uk/projects/fastqc/) to determine the quality of the raw reads, followed by RSeQC (v3.0.1) [19] that was used to determine the alignment quality and distribution of reads over the reference genome (number of mapped reads, proportion of uniquely mapped reads). Finally, the software MultiQC (v0.9) [20] was used to aggregate all QC results across samples.

A total of 126 samples were sequenced using 2×75bp or 2×100bp pair-end reads (**Figure 2A**). On average, a total of 64±7.4 paired-end million reads were generated with a per sequence mean quality score higher than 35 (**Figure 2A**). Given the high coverage and the lower length expected for a human transcriptome, the percentage of duplicated reads was high, reaching 69±5.5%, but the proportion of overrepresented sequences was low (<2%) indicating that all RNA sequenced libraries were diverse. The report of STAR alignments showed that on average 96.8±1.2% of the reads mapped to the reference genome with 91±2.3% mapping to unique loci (**Figure 2B**). The 3.15±1.26% of unmapped reads correspond mainly to reads with a short-alignment length (3.05±1.24%) that might result from the trimming process (trim of adaptor or low-quality bases, **Figure 2B**). Finally, as expected most of the MESOMICS reads mapped to mRNA structures including CDS and UTR regions (86.2±3.1%, **Figure 2C**).

**Figure 2.**
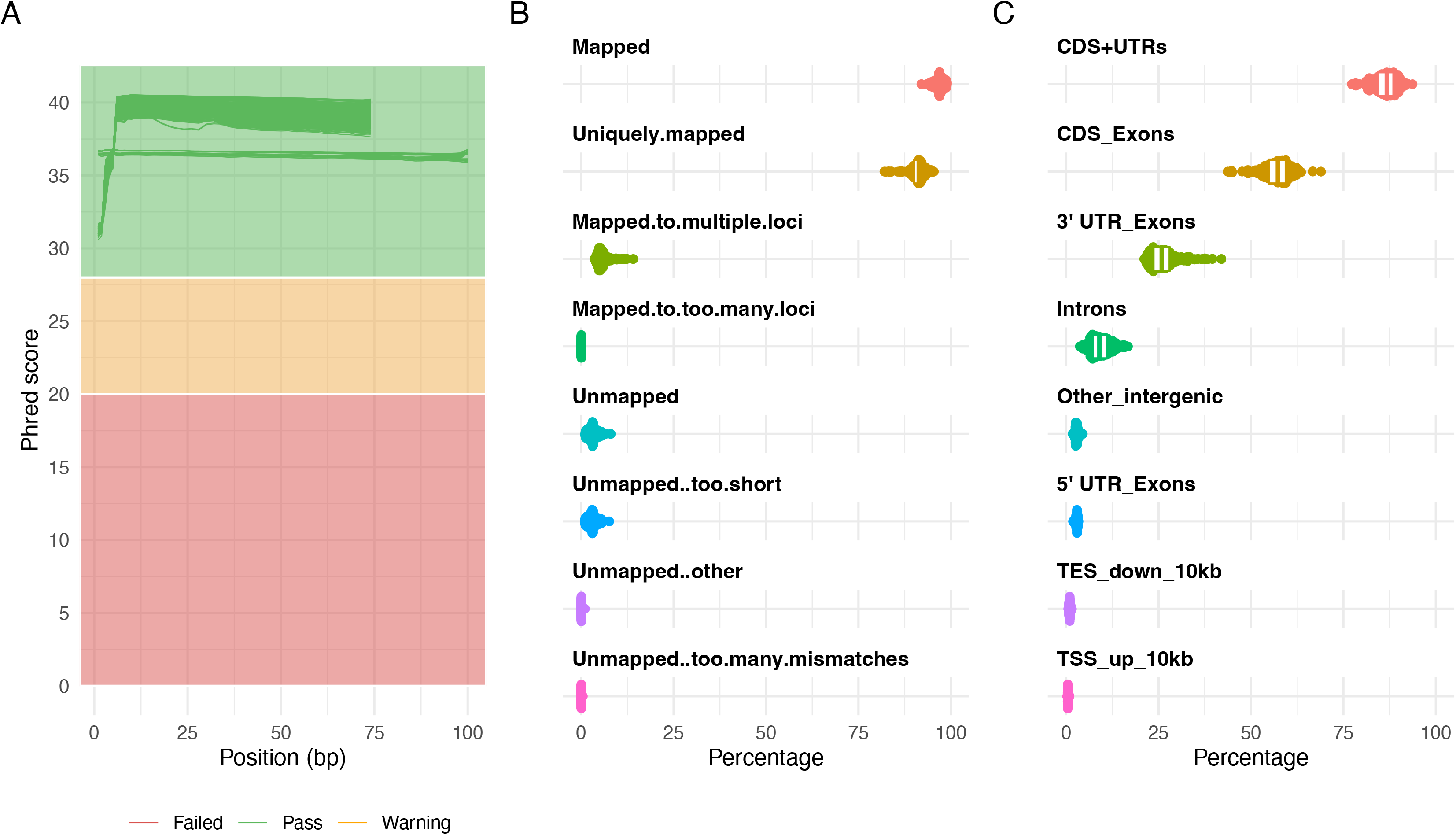
Quality control of RNA Sequencing (RNA-seq) data. A) Distribution of sequence quality scores in Phred scale for 2×75bp and 2×100bp read pairs. B) STAR alignment scores. C) Distribution of reads mapped to different genomic regions.

#### DNA methylation data

DNA methylation analyses were performed in-house for 135 MPM samples from 122 patients, and an additional two technical replicates and three adjacent normal tissues, with Infinium EPIC DNA methylation beadchip platform (Illumina) which interrogates over 850,000 CpG sites, as described in [13]. Resulting raw IDAT files were processed using our in-house workflow (https://github.com/IARCbioinfo/Methylation_analysis_scripts, commit SHA bcfe876) in the R statistical programing environment using R packages minfi (v1.34.0) and ENmix (v1.25.1), and consisted of the following four steps: pre-processing quality control, functional normalization, probe filtering, and finally beta and M-value computation.

During quality control checks on the raw data, one poor quality sample was identified when comparing per sample log2 methylated and unmethylated chip-wise median signal intensity (function getQC, minfi, **Figure 3A**) which was subsequently removed, and all samples displayed an overall *p*-detection value < 0.01 (function detectionP, minfi). Functional normalization, probe filtering, and beta and M-value computation were performed as described in [13]. The resulting dataset consisted of beta and M-values for 139 samples across 781,245 probes, with the M-value table containing nine -∞ values which were replaced by the next-lowest M-value for statistical analysis. The effect of normalization and probe-removal on DNA methylation profile is shown as beta density plots (pre-normalization in **Figure 3B** and post-normalization and probe removal in **Figure 3C**). Principal components analysis (PCA) was performed to detect batch effects, and to examine the effect of normalization, this was performed on a reduced number of samples (*n*=122, one tumor per patient, excluding technical replicates and normal tissues). Two datasets were used: (i) pre-normalized, unfiltered M-values (obtained from the GenomicRatioSet, function getM, minfi), and (ii) normalized and filtered M-values. Dataset (i) contained 2,478 CpGs with at least one NA value which were omitted before PCA, and 21,969 -∞ values were replaced with the next lowest M-value in the dataset, leaving an M-value matrix of 863,381 probes.

**Figure 3.**
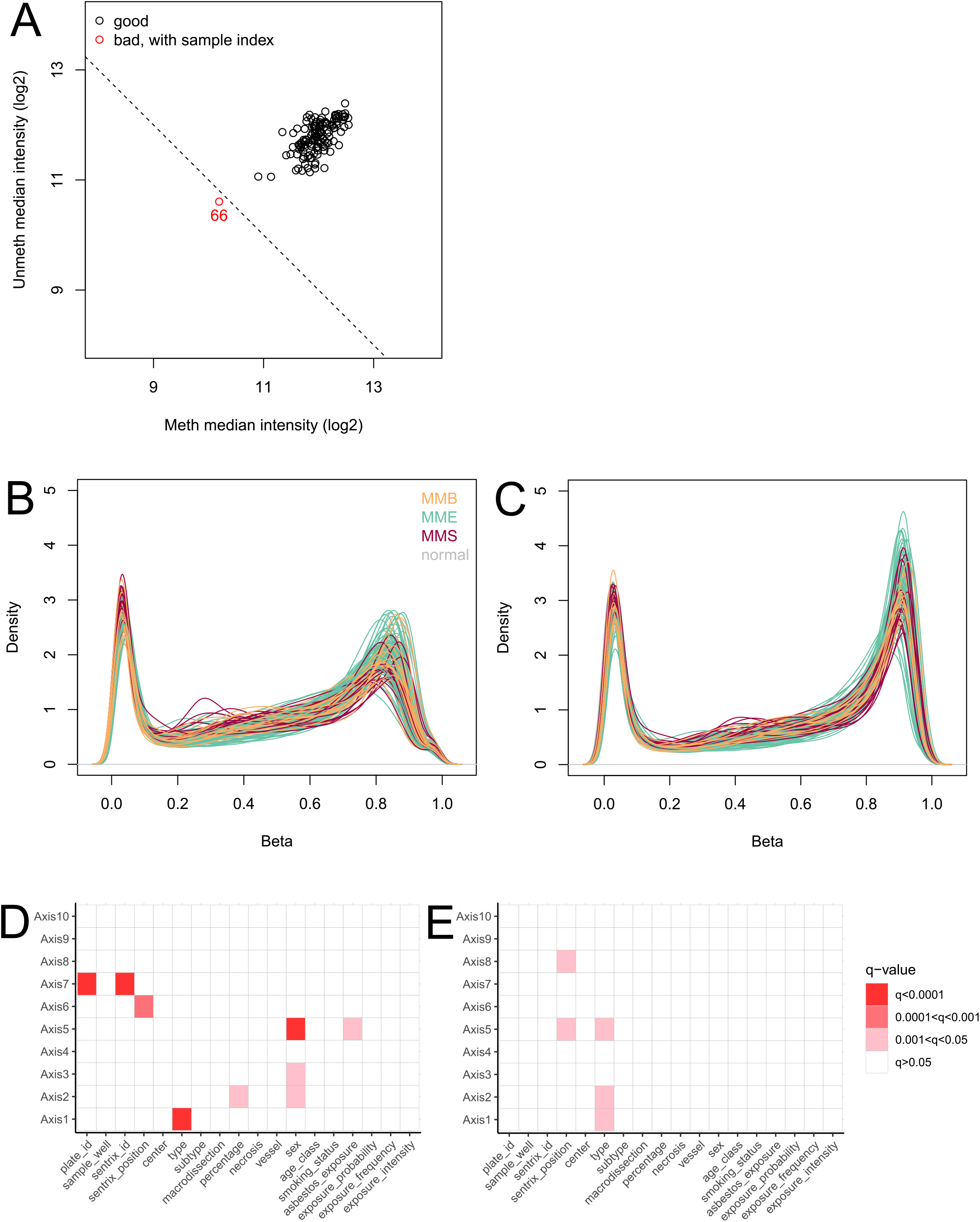
Quality control of EPIC array sequencing data. A) Signal intensity plot. Log2 methylated and unmethylated median signal intensity plot of 140 samples. One sample (coloured red) fell below the cut-off of 10.5 and was subsequently removed from analysis. B) Pre-normalisation beta density plot. Beta density plot of 140 samples across 865,859 probes, coloured by tumour/normal type, prior to functional normalisation. C) Post-normalisation and filtering beta density plot. Beta density plot of 139 samples across 781,245 probes, coloured by tumour/normal type, following functional normalisation and removal of cross-reactive, sex-chromosome, SNP, and failed (p-detection > 0.01) probes. D) Association of technical and clinical variables with pre-normalisation principal components. Association of technical and clinical variables with principal components one to ten, for 122 samples. Principal components calculated from M-values of 863,381 pre-normalised probes. E) Association of technical and clinical variables with post-normalisation principal components. Principal components calculated from M-values of 781,245 probes following functional normalization and probe removal.

R package ade4 (v1.7-15) was used to calculate the first 10 principal components (function dudi.pca) across each dataset individually. We checked the association of the first 10 PCs with technical (chip, position on the chip, batch, sample well, sample provider, macrodissection), clinical (sex, age class, and smoking status), morphological (histopathological type, subtype, tumor percentage, necrosis and vessel level), and epidemiological variables (asbestos exposure, exposure probability, exposure frequency, and exposure intensity) using PC regression analysis, fitting separate linear models to each principal component with each of the 18 covariables of interest and adjusted the *p*-values for multiple testing (**Figures 3D** and **3E**). The first ten principal components in the normalized, filtered methylation data were significantly associated with type (PCs 1, 2, and 5), and sentrix chip position (PC 5, PC 8). The contribution of variance in the data from technical features before normalization was more pronounced, with sentrix chip and plate also being significant (PC 7), indicating functional normalization reduced technical batch effects on DNA methylation profile while retaining biological effects such as histological type. Before normalization, sex was significantly associated with PCs 2, 3, and 5, but not associated with any PCs in the normalized, filtered dataset. As probes on the sex chromosomes were removed after normalization, it was expected that this would reduce the effect of sex on variance in the dataset.

### WGS variant calling in tumor-only samples

#### Copy number variants

Somatic Copy Number Alterations (SCNA) were called using our nextflow workflow purple-nf (https://github.com/IARCbioinfo/purple-nf, v1.0) that implements the PURPLE [21,22] software for matched and tumor-only WGS samples. To assess the quality of tumor-only PURPLE calls, a total of 57 matched-pairs were used as an evaluation set. Briefly, we ran PURPLE twice for each matched sample (**Figure 4A**): first using as input the matched-pairs, and second using only the tumor WGS as input. Subsequently we performed a direct comparison of the PURPLE tumor-only calls with their corresponding matched-pair calls for the following features: tumor purity, ploidy, number of segments, percentage of diploid, amplified, and deleted genome regions, as well as major and minor copy number states at the gene-level (**Figure 4**).

**Figure 4.**
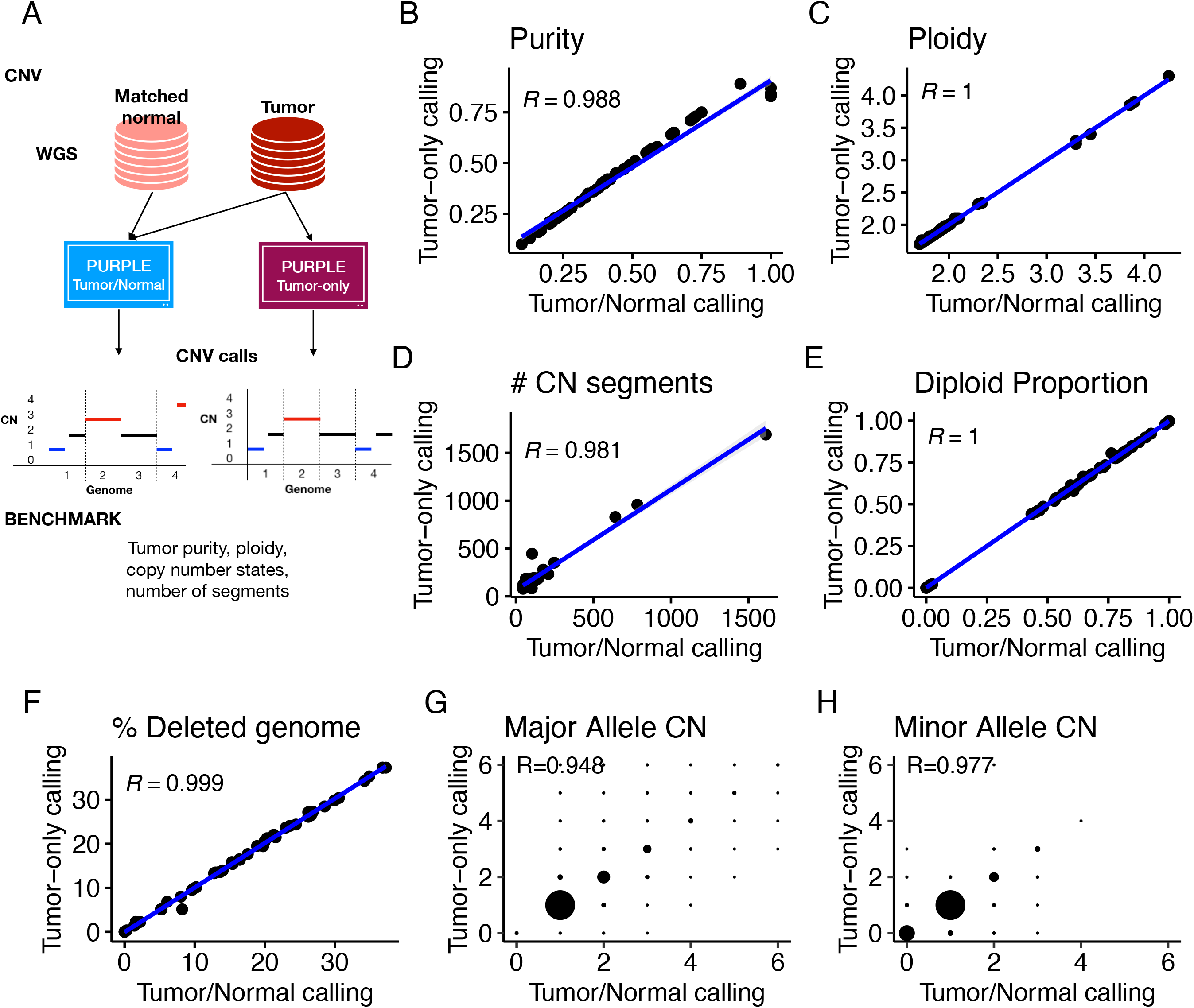
Performance of somatic copy number variant calling from tumor-only samples. A) Schematic of the benchmarking procedure. Comparison of Tumor/Normal and Tumor-only calling for B) Purity, C) Ploidy, D) Number of copy number segments, E) Diploid proportion, F) Percentage of deleted genome, G) Major allele copy number and H) Minor allele copy number.

This benchmarking revealed a high concordance across all the evaluated metrics between tumor-only and matched PURPLE calls. Indeed, the agreement for purity (**Figure 4B**, *R* = 0.988), ploidy (**Figure 4C**, *R*=1), number of copy number segments per tumor (**Figure 4D**, *R* = 0.981), and percentage of genome changed (diploid, amplified, and deleted) exceeded a 0.98 correlation (**Figure 4 E-F**). Moreover, a high concordance was also observed at the gene level with major and minor copy number alleles reaching *R* > 0.94 (**Figure 4 G-H**). Finally, the only detected issue of tumor-only calls was observed near telomeric and centromeric regions, where artefactual focal peaks were detected (**Supplementary Figure 1**). These problematic regions were manually curated and the copy number segments overlapping such regions were removed from the tumor-only calls (see list of excluded segments in Table S1). In addition, because PURPLE does not round copy number values to 0, but rather penalizes negative values in the model fit, for all samples (both matched and tumor-only), following similar discussions with the PURPLE developers on the handling of negative values (https://github.com/hartwigmedical/hmftools/issues/102), we rounded slightly negative copy number estimates (in]-0.5,0[) to 0 and excluded largely negative copy number estimates (<-0.5) from subsequent analyses, because they suggest high noise in the read depth and are thus unreliable calls. Note that in total (including segments with largely negative values), we excluded only 0.26% of the total segment length.

#### Point Mutations

Unlike copy number variants whereby the software (PURPLE) directly generated highly accurate results in tumor-only mode without any post-processing, for point mutations direct outputs from the software (Mutect2, [23]) and typical filters (i.e. removing variants matching germline databases) did not remove at high accuracy the germline variants present in tumor-only WGS. Therefore, we trained and evaluated the performance of a supervised machine learning model based on a random forest (RF, [24]) for distinguishing germline from somatic variants in tumor-only WGS (**Figure 5A**).

**Figure 5.**
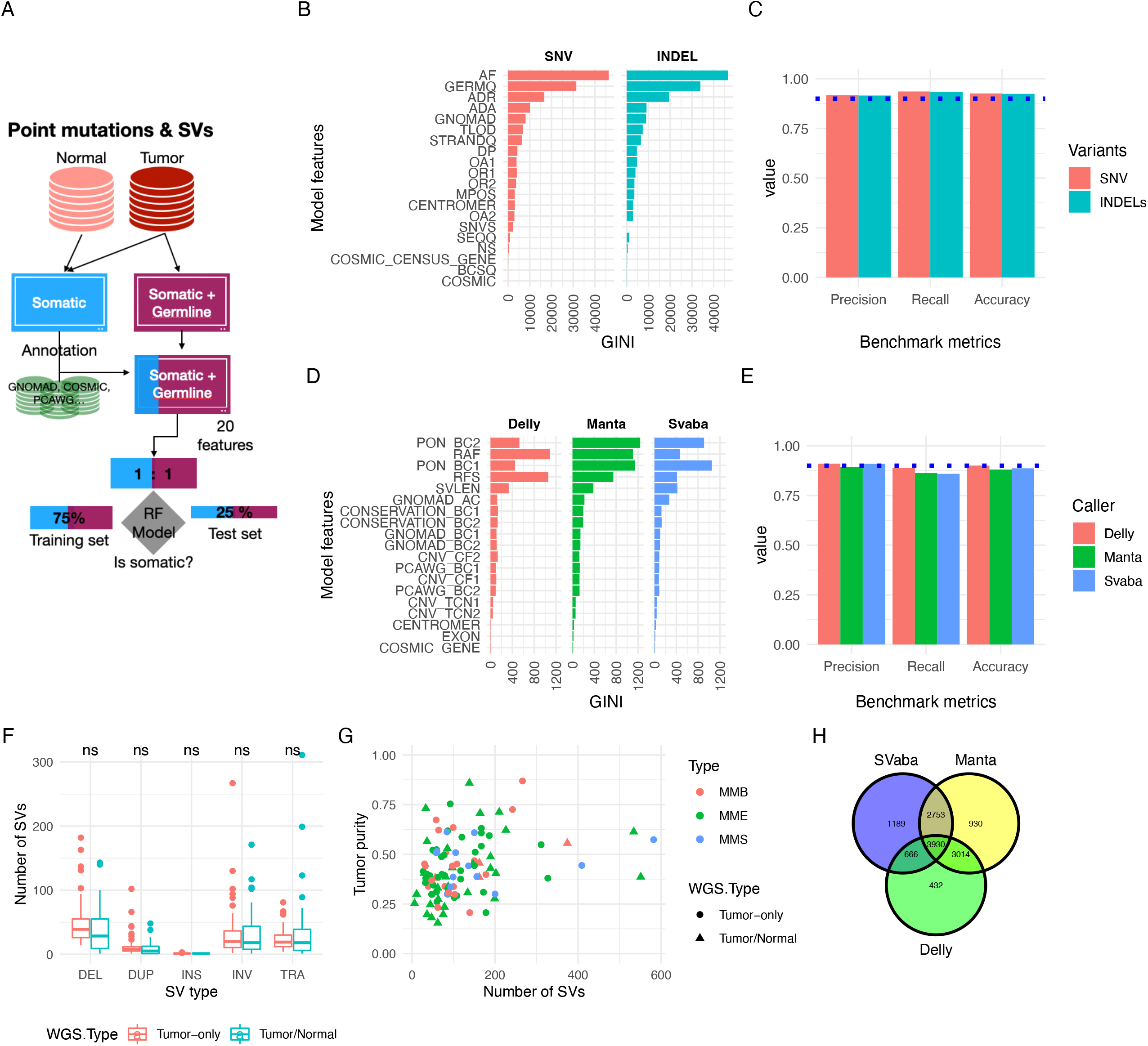
Performance of somatic point mutation and structural variant calling from tumor-only samples. A) Schematic of the benchmarking procedure. B) Random forest model features and their ranking for predicting somatic SNV and Indels. C) Performance metrics (precision, recall, accuracy) for classifying somatic point mutations with the best performing RF models. D) SVs Random Forest model features and their ranking for predicting somatic SVs. E) Performance metrics for classifying somatic SVs. F) Number of SVs as function of WGS type. Mean comparison between WGS types was performed using a t-test with no significant (ns) result found. G) Number of SVs as function of tumor purity. A linear model (number_sv ~Purity*WGS_type*SubType) was built to predict the number of SVs, no significant coefficients (p.value < 0.05) were found. H) Venn diagram of the final consensus MESOMIC SVs set.

The matched samples were used as input for training and evaluating the performance of the random forest (RF) model (**Figure 5A**). Point mutations were called using Mutect2 using our nextflow pipeline Mutect2-nf (https://github.com/IARCbioinfo/mutect-nf, v2.2b). To build the RF model, we chose a total of 20 features divided into three main classes, namely: associated with external databases, genomic location/impact, and features obtained directly from the Mutect2 variant caller (**Figure 5B**). For the first class of features, we used the gnomAD (r3.0 with 526M SNPs and 69M indels) [25] and COSMIC (v90, with 18.6M SNVs and 0.99M indels) [26] databases as reference for germline and somatic variants, respectively. The GNOMAD feature encodes the minor allelic frequency of each variant matching a variant present in the gnomAD database, otherwise providing a value of 0. The matching of variants was performed using bcftools (v1.10.2, annotate function) [27]. The COSMIC and COSMIC_GENE_CENSUS features are Boolean variables encoding if the variant matched one present in the COSMIC database or if its genomic location overlaps with the boundaries of a COSMIC gene (*n*=723). The second class of features included substitution patterns, functional impact, and genomic location of variants. The substitution patterns (i.e. mutation C->A) were converted into six categorical values (SNVS model feature), as somatic variants are often enriched in particular signatures [28]. The BCSQ feature encodes the impact of variants into four categories (MODIFIER, LOW, MODERATE, and HIGH) according to how the mutation affects coding genes. Variant impact was calculated using the bcq [29] function of bcftools and the classification of impact was performed with the Ensembl variant consequences table (https://www.ensembl.org/info/genome/variation/prediction/predicted_data.html). The CENTROMERE feature indicates if a variant is located in centromeric regions. Finally, fourteen features were derived from the Mutect2 caller, which are associated with read depth and orientation (DP, ADR, ADA, OR1, OR2, OA1, and OA2), sequencing errors and artifacts (MPOS, STRANDQ, SEQQ, TLOD, GERMQ), frequency across samples (NS), and allele frequency (AF, **Figure 5B**).

For training the RF model a total of 46 tumors with matched normal MPM whole-genome sequences called with both the tumor-only and matched modes of Mutect2 were used (**Figure 5A**). The matched somatic calls (ground-truth) were used to annotate the variants of the tumor-only WGS into germline and somatic classes. The number of germline variants called in tumor-only mode (*n*=694,273) greatly exceeded the number of somatic variants (*n*=203,993) generating an imbalance between classes. Therefore, we subsampled the germline class as a function of the somatic class to mitigate bias arising from class imbalance during training (1:1 somatic:germline ratio, *n*=407,984). To determine optimal values for the RF parameters we performed a grid search for tuning the mtry (4, 8, 12, 16), ntree (500, 1000, 1500), and nodesize (5, 25, 50, 100) parameters in a total of 48 RF-models. The training and evaluation of models was performed using 75% and the remaining 25% of the dataset, respectively. The grid search revealed that the optimum parameters were mtry=8, ntree=1000, and nodesize=5, reaching a model accuracy of 0.9276 in the testing set. Additionally, the minimum accuracy reached by any RF model was 0.9220, indicating that the parameter optimization had a marginal impact on the RF models. A random forest model for SNVs (rfvs01) was trained with the optimum parameters using a total of 326,388 (80%) variants (1:1 ratio). Analysis of the feature importance revealed that the allele frequency (AF) is the most discriminative feature included in the model (**Figure 5B**). For indels, a random forest model (rfvi01) was built with the same optimal parameters using a total of 337,442 variants (1:1 ratio, including 305,988 SNVs and 31,454 indels) and removing the SNVs feature. The performance of the optimal RF-models for SNVs and indels reached an accuracy of 0.926 and 0.924, respectively (**Figure 5C**). To further control the false positive rate (overall FDR=6.4%) we used different cut-offs (RF probability) to classify as somatic coding (>0.5) and non-coding (>0.75) variants. Finally, the trained RF models (rfvs01 and rfvi01) were used to classify a total of 1,454,942 variants (SNVs=1,317,200 and indels=137,742) of which 217,436 variants (including SNVs and indels) were classified as somatic. With these results we have developed a highly accurate and robust methodology to call SNVs and indels in tumor-only WGS datasets for which a series of matched tumor-normal samples are available. The source code and the random forest models implemented are available in the Github repository at https://github.com/IARCbioinfo/RF-mut-f.

#### Structural Variants

Large genomic rearrangements were detected using a consensus variant calling approach including SvABA (v1.1.0)[30], Manta (v1.6.0)[31], and Delly (v0.8.3)[32] followed by subsequent integration with SURVIVOR (v1.0.7)[33]. Our nextflow pipeline implementing the consensus variant calling approach for matched WGS is available at https://github.com/IARCbioinfo/sv_somatic_cns. In brief, each structural variant caller was run following the recommended practices (filters and excluding masked genomic regions). Consensus between callers was built using SURVIVOR (merge command) by computing an all-versus-all structural variants (SVs) comparison. Matching pass-filter somatic SV pairs were considered when both breakpoints overlap at a maximum distance of 1kb. Somatic SVs called by at least two callers as well as single-tool SVs supported by > 15 pair-end reads were included in the consensus set.

Like for point mutations, we implemented custom random forest models to distinguish at high accuracy somatic from germline SVs in tumor-only MESOMICS samples (**Figure 5A**). The RF-models were composed of a total of 19 features based on external databases, a custom panel of normal, genomic regions, and SV features obtained directly from each SV caller (**Figure 5D**). The gnomAD database (v2.1, *n*=299,211) GRCh38 liftover (dbVar https://ftp.ncbi.nlm.nih.gov/pub/dbVar/data/Homo_sapiens/by_study/vcf/nstd166.GRCh38.variant_call.vcf.gz) was used as reference for germline SVs. Features describing the frequency of the SV in gnomAD (GNOMAD_AC) and the number of germline SVs around each breakpoint (10kb window, GNOMAD_BC1 and GNOMAD_BC2) were included in the RF SV model. The PCAWG consensus call set for SVs (v1.6, *n*=309,246) [34] was used as a reference of somatic SVs. The PCAWG SVs are in hg19 genome coordinates, thus we performed a liftover to GRCh38 using CrossMap (v0.3.9) [35]. PCAWG SVs at sample level (*n*=2,748) were merged into a non-redundant cohort callset with SURVIVOR (merge subcommand) leading to a total of 283,980 non-redundant somatic SVs. Two features associated with the number of somatic SVs around each breakpoint (10kb window, PCAWG_BC1 and PCAWG_BC2) were included in the model. To further enhance the germline filtering we generated a custom panel-of-normal (PON) for each SV caller using our set of normal samples (*n*=46). Atotal of 56,572, 52,915, and 24,381 non-redundant PON SVs were collected for Manta, Delly and SvABA, respectively. Two features associated with the number of PON SVs around each breakpoint (10kb window, PON_BC1 and PON_BC2) were included in the model. For genomic regions, we annotated the SV breakpoints with features associated to known cancer genes (COSMIC_GENE, cosmic v90), coding exons (EXON, gencode v33), centromeres (CENTROMER), and conserved genomic regions (100-way PhastCons) [36]. Finally, we included in the model features indicating the total copy number state (CNV_TCN1 and CNV_TCN2 for both breakpoints), tumor cell fraction estimations (CNV_CF1 and CNV_CF2 for both breakpoints), SV length (SVLEN), SV read depth (RFS), and SV alternative allele frequency (RAF). Our hypothesis behind all the aforementioned features is that the underlying genomic context of somatic SVs is different from those of germline SVs and that a random forest model should be able to distinguish both SV classes using the proposed features.

The training (75%) and evaluation (25%) of the random forest model for each SV caller was performed using a total of 12,454, 16,720, and 12,264 SVs at 1:1 somatic:germline proportions for Delly, Manta, and SvABA, respectively. All three SV random forest models were trained using the default random forest parameters (mtry=4, ntree=400 and nodesize=1). The precision, recall, and accuracy achieved by each model were 0.905±0.009, 0.87±0.016, and 0.889±0.010, respectively (**Figure 5E**). The most important features of the models were the number of PON SVs around both breakpoints, SV alternative allele frequency, SV read depth, and SV length (**Figure 5D**). To control the FDR, SVs located in coding and non-coding genomic regions were classified as somatic if their random-forest probability exceeded 0.5 and 0.75, respectively. Additionally, somatic SVs matching SVs present in the MESOMICS PON or located in centromeric regions were discarded. A consensus of SV sets for each tumor-only sample was built using the same steps performed for the matched WGS. Finally, SVs having a frequency higher than four across all the tumor-only samples were also classified as potentially germline and removed from the final consensus set. We performed additional comparisons by SV type, SV length, and number of SVs as a function of the purity of samples, WGS type(Tumor/Normal, Tumor-only) and MPM subtype and did not observe any significant difference between SVs called in the tumor-only or matched WGS MESOMICS series (**Figure 5F** and **5G**).

The SV calls for the MESOMICS tumor-only samples include a total of *n*=8,229 SVs, which combined with the SVs called in the matched series gave a total of *n*=12,914 (**Figure 5H**). With these results we have developed a highly accurate and robust methodology to call SVs in tumor-only WGS datasets for which a series of matched tumor-normal samples are available. The source code and the SV random forest models are available in the Github repository at https://github.com/IARCbioinfo/ssvht.

### Data Validation

#### Muti-omic sample matching

The software NGSCheckMate was used to check the match between sequencing modalities of a given MESOMICS patient. NGSCheckMate was run using our nextflow implementation available at https://github.com/IARCbioinfo/NGSCheckMate-nf (v1.1a). NGSCheckMate, using WGS and RNA-seq, confirmed that the majority of MESOMICS samples were correctly paired (**Figure 6A**, black segments). However, NGSCheckMate discovered that the WGS of MESO_094_T and MESO_096_T matched (**Figure 6A**, red segments). Further examination of these samples confirmed that both WGS come from the same patient but were annotated differently during sample collection. In addition, the RNA-seq replicate named MESO_054_TR1 matched with the group of samples coming from patient MESO_051. After sequencing a second RNA-seq aliquot from MESO_054_T, named MESO_054_TR2, and re-performing the NGSCheckMate analysis, we confirmed a miss-annotation of these RNA-seq samples and proceeded to rename them as MESO_051_TR1 and MESO_051_TR2, respectively. After the aforementioned corrections, all the sequencing modalities at the sample and patient level were correctly paired for the complete MESOMICS cohort.

**Figure 6.**
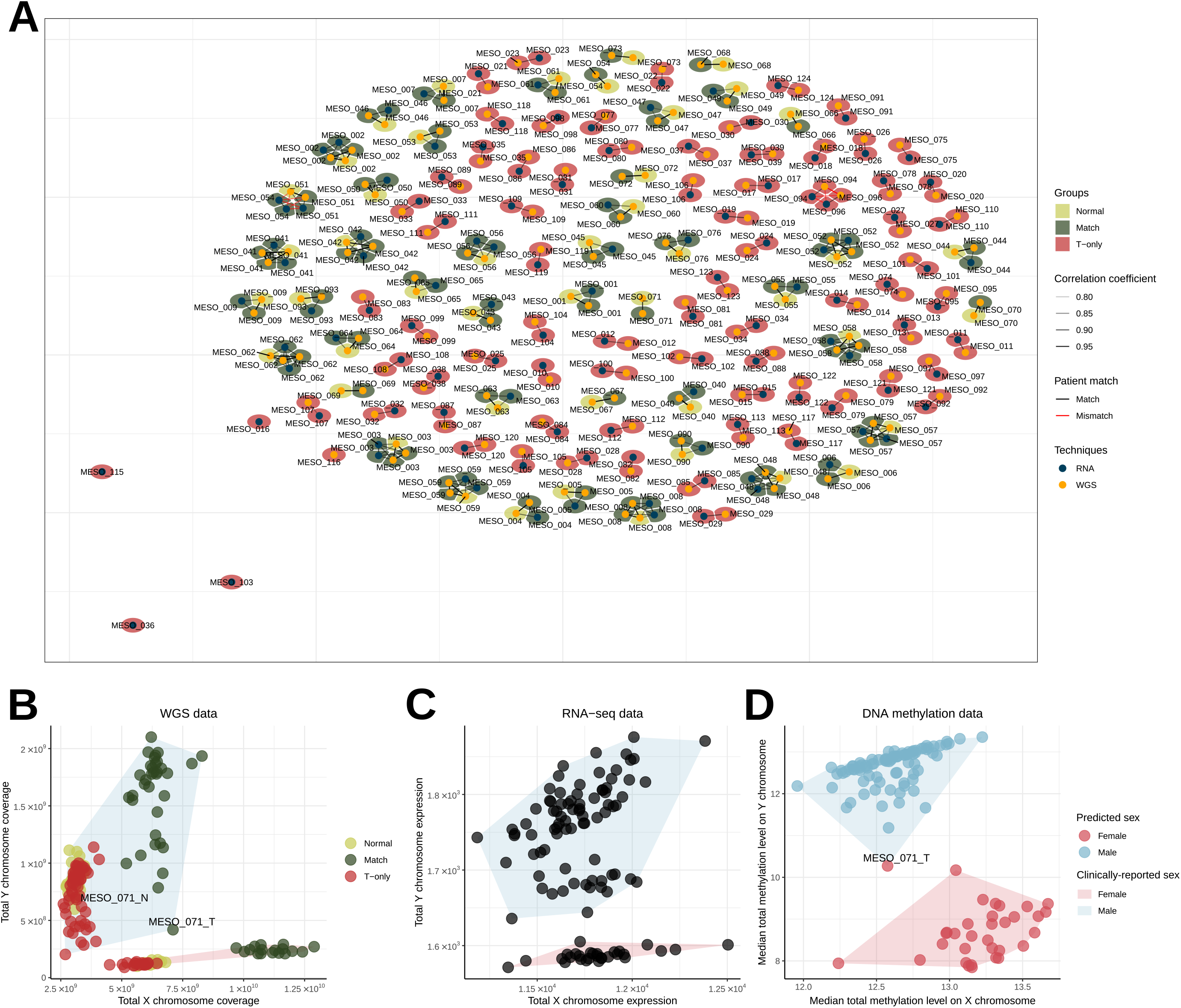
Applications of data validation using multi-omics data A) Network of matching WGS and RNA-seq samples, as computed by software NGSCheckmate. Edge transparency corresponds to the Pearson Correlation r between SNP panel allelic fractions; node color and surrounding color correspond respectively to the techniques (WGS or RNA-seq) and to the tissue type (Normal, Matched samples or T-only samples). B-D Sex reclassification and multi-omic validation of reported clinical sex. B) Total exome reads coverage on the X and Y chromosomes for each sample. C) Total expression level of each sample on the X and Y chromosomes (in variance-stabilized read counts). D) Median methylation array total intensity on the X and Y chromosomes. In panel (B), point colors correspond to the WGS groups: normal samples in light green, tumor samples with matched normals (Match) in dark green, and tumor samples without matched normal (T-only) in red. In each panel, filled polygons correspond to the sexes given by the clinical annotations (blue for male, red for female). In panel D) point colors correspond to the sexes predicted by the DNA methylation QC. Samples with discordant reported clinical sex and molecular patterns on sex chromosomes are indicated.

#### Sex validation

We registered the sex (M for male or F for female) data for all the 124 patients of the MESOMICS cohort. We validated the sex annotation based on the concordance of whole-genome, transcriptome, and methylome data (**Figure 6B-D**). First, the concordance between sex reported in the clinical data and WGS data was assessed by computing the total coverage on X and Y chromosomes (**Figure 6B**). Interestingly, some tumors from male individuals displayed an intermediate coverage on chromosome Y between other male and female cases, compatible with the large copy losses identified in our study like for example the tumor from MESO_071. Second, the concordance between sex reported in the clinical data and sex chromosome gene expression patterns (transcriptome) was performed by comparing the sum of variance-stabilized read counts (vst function from R package DESeq2, v.1.14.1) of each sample on the X and Y chromosomes (**Figure 6C**). Third, the concordance between the sex reported in the clinical data and the methylation data was assessed using a predictor based on the median total intensity on sex-chromosomes, with a cut-off of −2 log2 estimated copy number (function getSex from minfi, v.1.34.0, **Figure 6D**). The only sex discordance was observed in MESO_071 tumor sample due to somatic copy number losses in the Y chromosome, but the whole genome sequencing from matched blood confirmed that this patient was male (Figure 6B). In summary, the sex data of the MESOMICS cohort was validated using a multi-omic approach that confirmed the sex of all the MESOMICS samples.

### An integrative and interactive MPM phenotypic map

#### Task specialization analysis using Pareto

In order to integrate the MESOMICS multi-omic data and investigate the association between the detected genomic events in this new large genomic cohort and the observed MPM phenotypes, we firstly performed a multi-omic summary of MPM using MOFA [37] and secondly performed a task specialization analysis to identify MPMs with natural selection for specific cancer tasks (see [13]). We performed task specialization analyses using the well-established Pareto optimum theory (ParetoTI method) [10]. The Pareto front model has been fitted to different sets of samples using the ParetoTI R package (https://github.com/vitkl/ParetoTI, v0.1.13) on MOFA latent factors (LFs), restricted to LF1, LF2, LF3, and LF4 due to their association with survival and extreme phenotypes (see [13]). In brief, according to the theory a molecular map would take a particular shape (polyhedra) if a trade-off exists between several cancer tasks performed by the tumors. Using MOFA axes, we found a triangle (polyhedra with three vertices) corresponding to *k* =3 archetypes in the LF2-LF3 space. According to the Pareto optimum theory, this pattern results from natural selection for cancer tasks, with specialized tumors close to the vertices of the triangle (representing archetypes), and generalists in the center. We have also replicated the same analyses (MOFA and ParetoTI) on the previously published multi-omic studies from Bueno *et al*. [2] *(n=181)* and Hmeljak *et al*. [3] *(n=73)*. R scripts to prepare matrices for each omic layer, as well as scripts to run MOFA and the Pareto analysis for the three cohorts are available in the Github repository dedicated for this data note paper at https://github.com/IARCbioinfo/mesomics_data_note.

#### Biological interpretation of the MPM phenotypic map

We inferred each archetype’s phenotype by performing integrative gene set enrichment analysis on the expression data and identified the following cancer tasks and tumor phenotypes: Cell division, Tumor-immune-interaction, and Acinar phenotype (see [13]). Tumors specialized in the Cell division task displayed upregulation of pathways within the “cell division” task as reported by Hausser et al. [38] in multiple tumor types. This phenotype was enriched for non-epithelioid tumors and presented higher levels of necrosis, higher grade, high expression of hypoxia response pathways, and greater percentage of infiltrating neutrophils that are innate immune response cells. Cell division specialization was supported by the high expression levels of the proliferation marker *MKI67*, and increased genomic instability. Tumors specialized in the Tumor-immune-interaction task carried upregulated immune-related pathways, high expression of immune checkpoint genes, and high immune infiltration with an enrichment for adaptive-response cells: lymphocytes B, T-CD8+, and T-reg. The last extreme phenotype was characterized by samples with Acinar morphology, presenting a very structured tissue organization with epithelial cells tightly linked into tubular structures, and correlated with the presence of monocytes and NK cells (innate immune response cells). This phenotype presented the lowest epithelial-mesenchymal transition (EMT) score [39], with overexpression of epithelial markers such as cell-adhesion molecules, corroborating the importance of tissue organization in this phenotype, and also low levels of *MKI67* expression, indicating slow growth. Altogether these data provide a biological understanding for the molecular and phenotypic heterogeneity characteristic of MPM tumors.

#### Reuse potential

The MESOMICS project represents the most comprehensive molecular characterization of MPM to date, made possible by inclusion of the largest WGS dataset yet reported, and by the depth of the analyses undertaken. Multi-omics integration and biological interpretation through the lense of Pareto theory has allowed us to uncover three specialized MPM tumor profiles [13]. In order to replicate these findings while minimizing batch effects associated with bioinformatics data processing, we have accessed and reprocessed the raw data from previously published MPM multi-omics studies [2,3] using the same analytical procedures. A by-product of this laborious work is the creation of the largest (*n=374* samples in total) existing harmonized dataset of MPM multi-omics data.

In order to maximize the reuse potential of this dataset, we have also harmonized the available clinical, epidemiological and morphological data from these three cohorts. In addition to providing the raw data, the full list of genomic variants and the entire matrices of expression and methylation levels, we provide a curated and harmonized list of molecular features (e.g. immune cell composition, measures of genomic instability, presence of whole genome duplication, copy number in recurrently altered regions, driver genes mutational status, expression level of some relevant genes etc.) across all samples (**Supplementary Table S2**).

This MPM phenotypic map has been shared on the TumorMap web portal [11], offering an interactive visualization of this data in the tumor phenotypes space (Cell division, Tumor-immune-interaction, and Acinar phenotype), including all the harmonized clinical, morphological, epidemiological, and molecular data attributes mentioned above. The TumorMap interface provides an interactive way to explore and navigate through the map, where each sample is represented by a dot localized according to its position in the phenotype space (**Figure 7**). The attributes can be used to change colors and filter samples, perform statistical tests, and new attributes can be derived from pre-existing ones using set operations. This flexible and user-friendly interface will enable new hypotheses to be tested without computational expertise, and expands the reuse potential of the dataset. The TumorMap is available at https://tumormap.ucsc.edu/?bookmark=746c4bc0e8bc4eb5f280cdd81c7dcc783955faf2e2b493d0d205b7d1e92b98c4.

**Figure 7.**
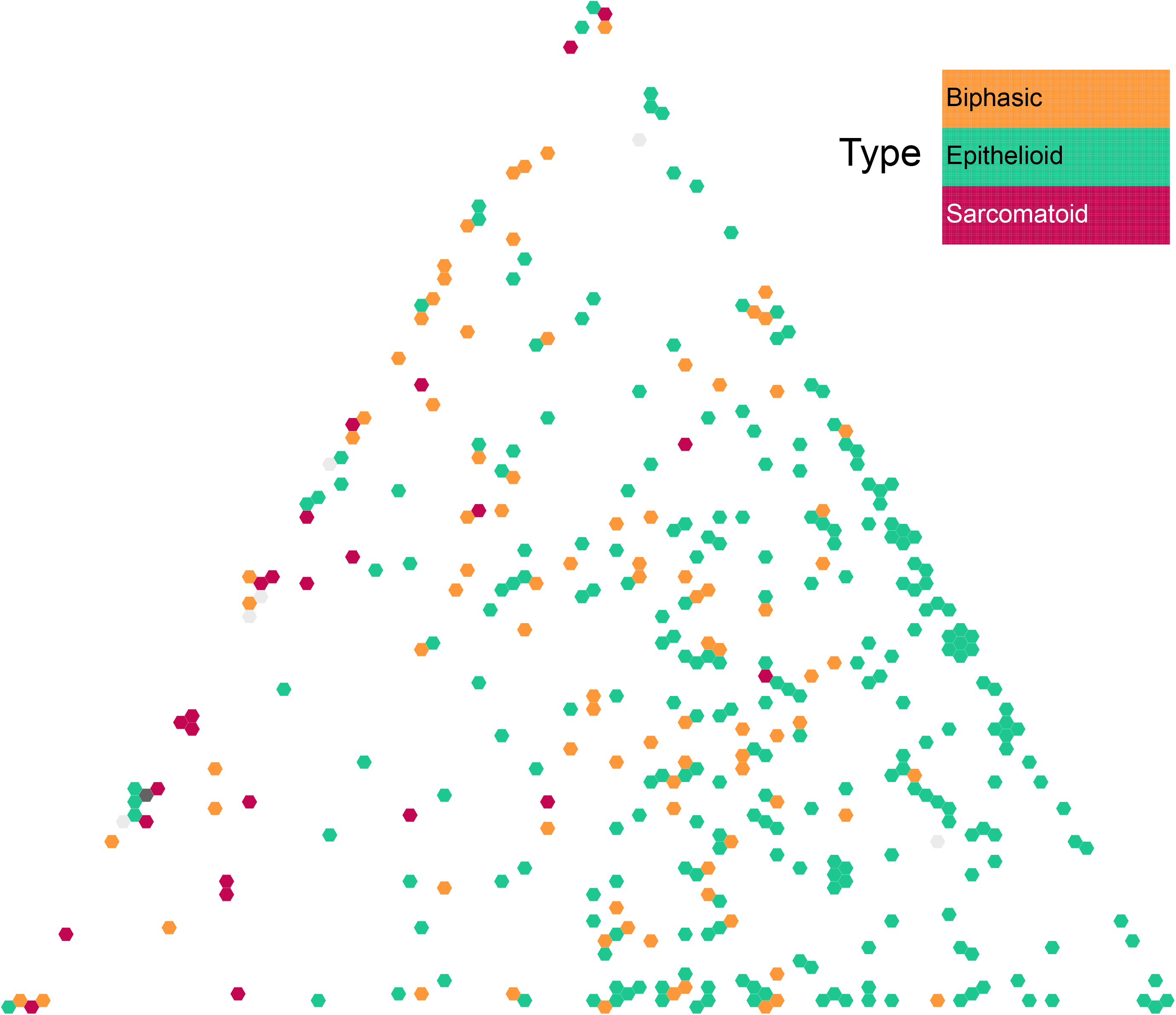
MPM molecular phenotypic map. Screen capture from the TumorMap portal, using the hexagonal grid view, each point representing a MPM sample in the triangular phenotypic space: cell division (left vertice), tumor-immune-interaction (top vertice), and acinar phenotype (right vertice). Point colors correspond to the histological types and can be interactively changed by the users on the web portal.

### Conclusion

We demonstrated that we provide a high-quality multi-omic dataset of malignant pleural mesothelioma, including the largest whole-genome sequencing dataset of malignant pleural mesothelioma to date, consisting of both raw and processed data, and important molecular phenotypes. By homogenizing the clinical, epidemiological, morphological and molecular data of our new series with the two previously published MPM multi-omics data series, we have created an unprecedented dataset for this rare cancer in terms of both size and detail. We provide all the resources to reproduce our analyses, as well as a user-friendly interactive visualization tool, which will contribute to advancing biological knowledge of this deadly disease.

## Availability of Supporting Data and Materials

The data used in this manuscript are available in the European Genome-phenome Archive (EGA), which is hosted at the EBI and the Centre for Genomic Regulation (CRG), under the accession number EGAS00001004812. R scripts are available in the github repository https://github.com/IARCbioinfo/MESOMICS_data. The interactive molecular phenotypic map is available at https://tumormap.ucsc.edu/?bookmark=746c4bc0e8bc4eb5f280cdd81c7dcc783955faf2e2b493d0d205b7d1e92b98c4.

## Supplemental files

**Supplementary Figure S1.** MPM CNV cohort profile aCNViewer plot [40] from Tumor-matched called as Tumor-Only (top), Tumor-only (middle), and Tumor-only after filtering (bottom). The circled regions correspond to artifactual peaks when calling CNVs with the Tumor-only mode of PURPLE. The aforementioned genomic regions were identified, filtered and are provided in Supplementary Table S1.

**Supplementary Table S1.** List of excluded genomic regions identified as artifactual when calling CNVs using PURPLE Tumor-only mode.

**Supplementary Table S2.** Harmonized and curated molecular, clinical, epidemiological and morphological data from our MESOMICS cohort, and the two previously published MPM multi-omics data [2,3]. This table can be explored interactively on the UCSC TumorMap web portal.

## List of abbreviations

ABRA: Assembly-Based Realigner
BAM: Binary Alignment Map
CDS: coding sequence
CNV: Copy Number Variant
CpG: cytosine–phosphate–guanine
EGA: European Genome-phenome Archive
EMBL-EBI: European Bioinformatics Institute
GATK: Genome Analysis Toolkit
IARC: International Agency for Research on Cancer
MME: malignant mesothelioma epithelioid
MMB: malignant mesothelioma biphasic
MMS: malignant mesothelioma sarcomatoid
MPM: malignant pleural mesothelioma
PCA: principal component analysis
QC: quality control
FR: Random Forest
RNA-Seq: RNA sequencing
SNP: single-nucleotide polymorphism
STAR: Spliced Transcripts Alignment to a Reference
TES: transcription end site
TSS: transcription start site
UCSC: University of California Santa Cruz
SV: Structural Variant
vst: variance-stabilized transformation
WGS: whole-genome sequencing

## Ethics approval

These data belong to the MESOMICS project, which has been approved by the IARC Ethical Committee.

## Author’s contributions

Conceptualization, M.F, N.A., L.F.-C; Methodology, M.F., N.A., L.F-C., L.M., A.DG., A.S.-O.; Software, A.DG., L.M., A.S.-O, C.V., N.A.; Validation, A.DG., L.M., A.S.-O, N.A.; Formal Analyses, A.DG., L.M., A.S.-O, C.V., N.A.; Investigation, A.DG., L.M., A.S.-O, C.V., N.A.; Data Curation, A.DG., L.M., A.S.-O, C.V., N.A.; Writing – Original Draft, A.DG., M.F, L.M., N.A., A.S.-O.; Writing – Review & Editing, A.DG., M. F., L.M., N.A., A.S.-O., L.F-C.; Visualization, A.DG, L.M., N.A., A.S.-O., C.V.; Supervision, L.F-C., M.F., N. A.; Project Administration, L.F.-C., M.F., L.M., N.A.; Funding Acquisition, L.F-C., M.F, N.A.

## Acknowledgements

This study is part of the MESOMICS project within the Rare Cancers Genomics initiative (www.rarecancersgenomics.com). We also acknowledge the Cologne Centre for Genomics (Cologne, Germany) and the Centre National de Recherche en Génomique Humaine (Evry, France) for generating high-quality sequencing data, as well as the Epigenomics and Mechanisms branch at IARC for generating high-quality methylation data. The results published here are part based upon data generated by the TCGA Research Network: https://www.cancer.gov/tcga.

