## Supplementary figures and images for "A molecular phenotypic map of Malignant Pleural Mesothelioma"

### Supplementary Figure 1

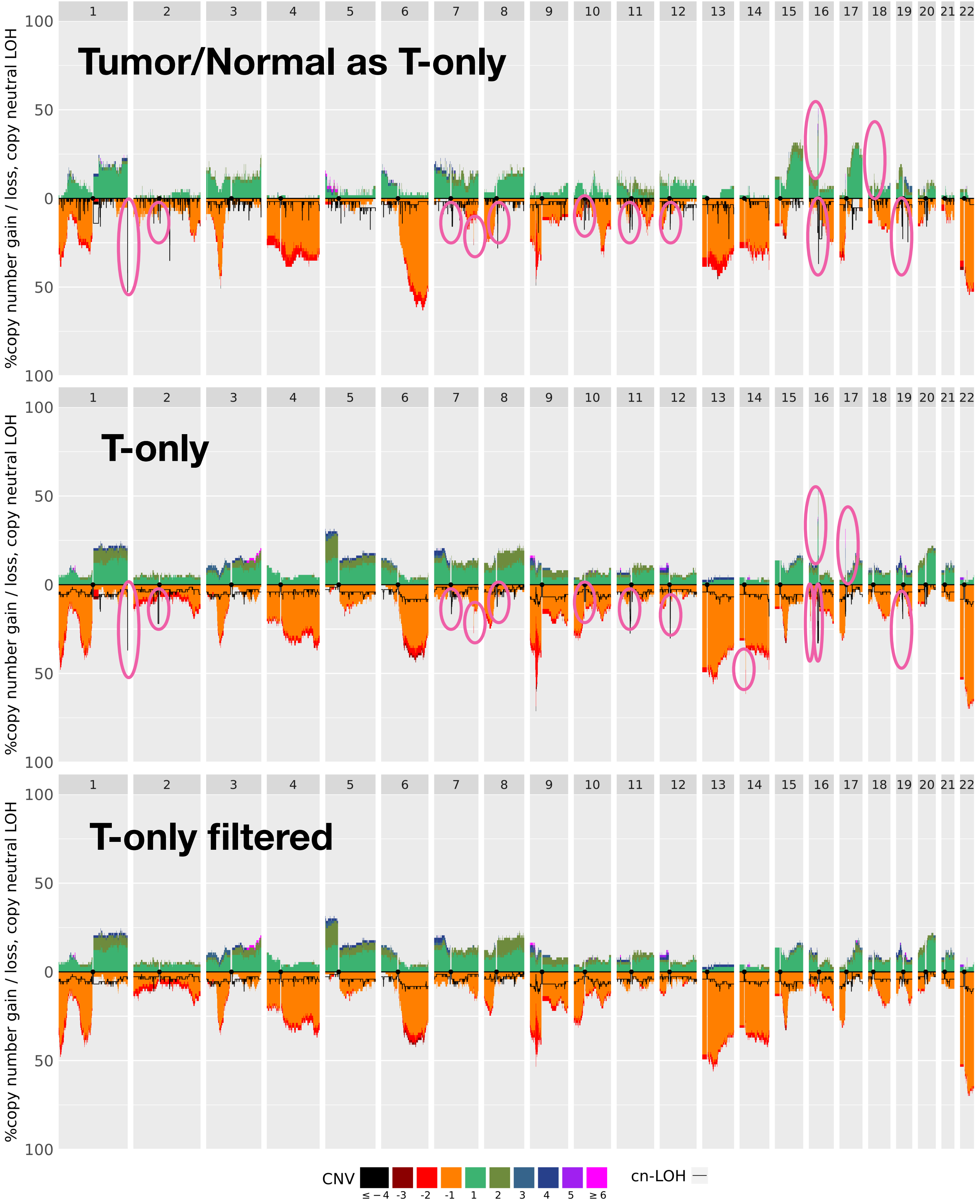
